# Taxonomy-aware, disorder-matched benchmarking of phase-separating protein predictors

**DOI:** 10.64898/2026.02.11.705241

**Authors:** Shuang Hou, Hexin Shen, Yong Zhang

**Author notes:** These authors contributed equally (Shuang Hou, Hexin Shen).

## Abstract

**Background:** Biomolecular condensates formed via liquid-liquid phase separation (LLPS) play vital roles in cellular organization and function. Computational prediction of phase-separating proteins (PSPs) is increasingly used to prioritize candidates at proteome scale, making robust, well-designed benchmarks essential for fair evaluation and iterative improvement of PSP predictors.

**Results:** We first show that a recently released PSP benchmark is substantially confounded by the imbalances in taxonomic origin and intrinsic-disorder compositions between positive and negative sets, allowing predictors to achieve high apparent performance by exploiting non-LLPS shortcuts and obscuring their true ability to distinguish PSPs. To minimize these effects, we construct a taxonomy-aware, disorder-matched PSP benchmark. Using this benchmark, we find that absolute sequence and biophysical feature values of PSPs differ markedly across taxa, whereas LLPS-associated feature shifts relative to taxon-specific proteome backgrounds are comparatively conserved. Benchmarking twenty PSP predictors under this framework reveals pronounced taxon-dependent variation in performance. Moreover, PSPs lacking IDRs consistently constitute a more challenging regime across methods, motivating routine disorder-stratified evaluation.

**Conclusions:** Our taxonomy-aware, disorder-matched benchmarking framework reduces shortcut-driven biases, enables more interpretable evaluation of PSP predictors, and provides guidance for developing models that capture transferable LLPS-associated signals rather than dataset- or taxon-specific shortcuts.

## Introduction

Biomolecular condensates are widespread membraneless cellular compartments that spatially and temporally organize biochemical reactions and have emerged as a pervasive principle of intracellular organization^1,2^. They support diverse biological functions ranging from transcriptional regulation to stress responses, and their dysregulation has been implicated in numerous diseases^3,4^. Condensates often form through liquid-liquid phase separation (LLPS), a demixing process in which macromolecules spontaneously concentrate into a dense phase while remaining dynamic and reversible. Proteins that participate in LLPS are commonly referred to as phase-separating proteins (PSPs). Given the diversity of PSP roles and the limited scalability of experimental characterization, a wide range of computational PSP predictors has been developed to infer LLPS propensity^5-24^. Accurate prediction is valuable for proteome-wide prioritization of candidate PSPs, but the credibility of these predictors depends critically on the datasets used for training and evaluation. Multiple LLPS-focused databases annotate proteins observed in biomolecular condensates, yet they differ substantially in conceptual scope and inclusion criteria, leading to highly divergent annotations and protein coverage. For instance, LLPSDB is oriented toward *in vitro* experimentally validated LLPS systems^25^; PhaSePro is restricted to experimentally validated driver proteins or driver regions^26^; PhaSepDB adopts broader inclusion by distinguishing PS-self proteins that phase separate without partners from PS-other proteins that require partners such as proteins or RNA^27^; DrLLPS organizes LLPS-associated proteins by functional roles as scaffolds, clients, or regulators^28^. These conceptual differences make benchmark construction non-trivial and underscore why carefully designed benchmarks are essential for reproducible and fair comparisons across predictors. Recently, the benchmark introduced by Pintado-Grima and colleagues^29^ represented an important step toward standardizing such comparisons by integrating curated positive and negative sets for systematic evaluation.

A key, often underappreciated complication is that the molecular grammars that encode condensate formation can vary across taxa. For example, yeast prions often share portable, intrinsically disordered prion domains enriched in asparagine, glutamine, tyrosine and glycine, whereas this domain architecture is not found in the mammalian prion protein or in the fungal prion HET-s and is rare in prokaryotes^30^. Such taxon dependence matters for benchmarking. As we show below, the Pintado-Grima benchmark couples PSP labels with taxonomic identity (with positives heavily enriched in mammals and negatives enriched in prokaryotes; see Results for details), and predictor performance on this benchmark is systematically associated with the similarity between a model’s training-taxonomic distribution and the benchmark’s composition. Under these conditions, benchmarks can reward taxon discrimination rather than LLPS-relevant biophysics, conflating genuine PSP generalization with differences in taxonomic composition.

A second axis of potential confounding arises from intrinsic disorder. Intrinsically disordered regions (IDRs) often contribute to multivalent interactions and are frequently associated with condensates, yet they are not universally necessary or sufficient for LLPS^31^. In parallel, LLPS can be driven by folded modular interaction domains that support multivalency through specific binding networks^31^. Therefore, a benchmark that inadvertently contrasts disorder-rich positives against disorder-poor negatives risks rewarding predictors that primarily detect disorder enrichment rather than LLPS-associated molecular grammars. This motivates controlling not only taxonomic composition, but also the balance of proteins with IDRs (IDPs) versus proteins without IDRs (non-IDPs) when constructing evaluation sets, in order to reduce shortcut learning.

Here, we address these issues by constructing a taxonomy-aware, disorder-matched PSP benchmark dataset designed to minimize the influence of taxonomic and disorder-related confounders. By balancing positives and negatives within major taxonomic groups and matching disorder composition between classes, our benchmark aims to prevent strong separation driven by taxonomic imbalance or disorder composition differences, thereby forcing evaluation to reflect LLPS-associated molecular grammars rather than trivial shortcuts. Using this benchmark, we show that absolute sequence and biophysical feature values of PSPs differ markedly across taxa, highlighting the importance of taxon-specific baselines. We therefore report PSP predictor performances separately for each major taxonomic group and further stratify by disorder status. Together, this framework enables more interpretable and trustworthy comparisons among PSP predictors and offers practical guidance for future model development.

## Results

### Pintado-Grima benchmark couples PSP labels with taxonomic identity

To determine whether evaluations on the PSP benchmark introduced by Pintado-Grima and colleagues are confounded by taxonomic composition, we first quantified the taxonomic-group distributions of its positive and negative sets. We partitioned proteins into seven major taxonomic groups: Animals(M) (mammals), Animals(NM) (non-mammalian animals), Fungi, Protists, Plants, Prokaryotes, and Viruses. This analysis revealed a pronounced imbalance between labels: positives were dominated by mammals (63.4%), whereas negatives were dominated by prokaryotes (51.7%), resulting in substantially different taxonomic profiles between the two classes (Fig. 1A). Thus, taxonomic origin is strongly coupled to the benchmark labels, raising the possibility that performance estimates on this benchmark can be inflated by taxon-associated signals rather than PSP-specific determinants.

**Figure 1.**
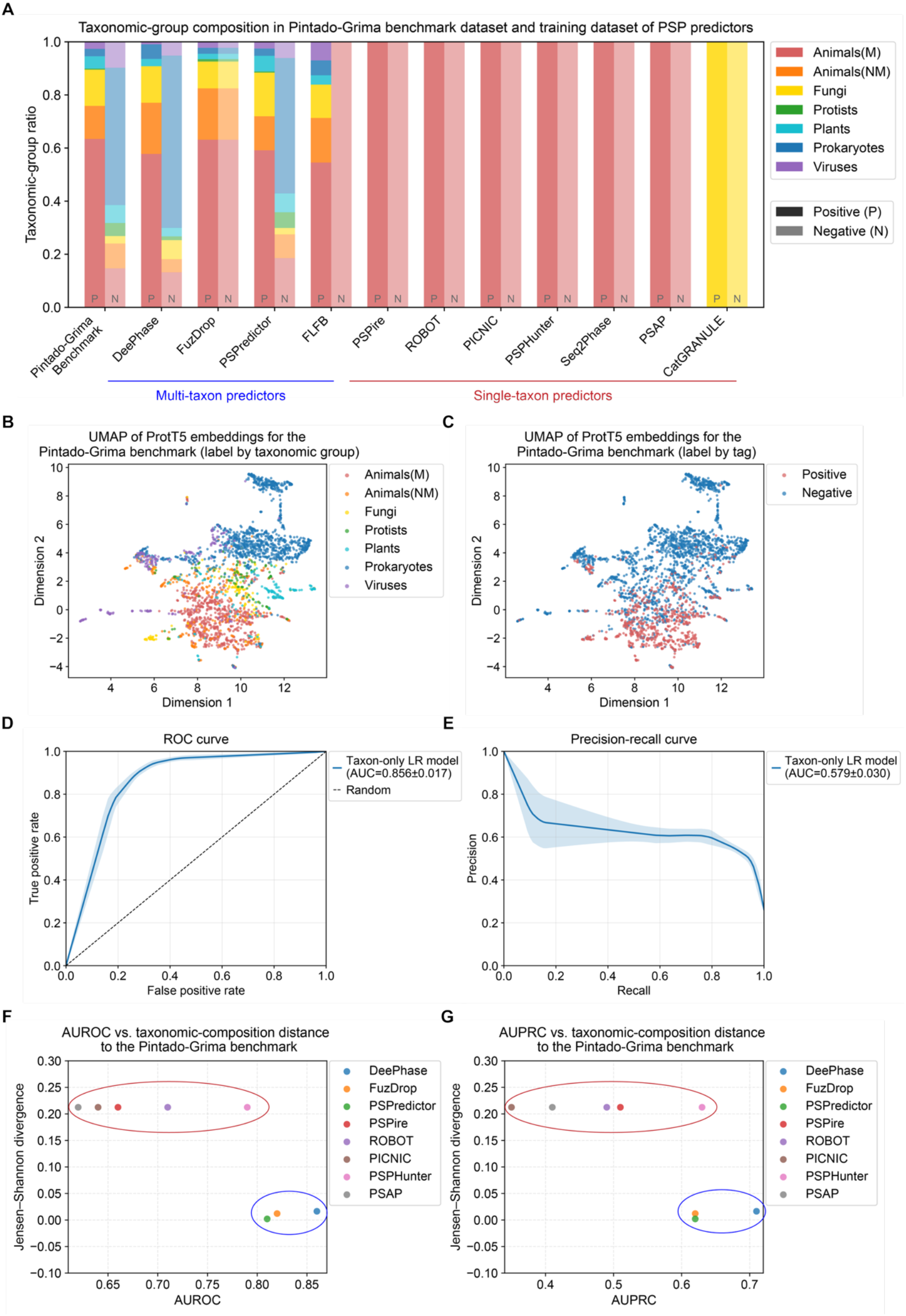
Benchmark performance is confounded by taxonomic composition in the Pintado-Grima dataset. **A** Taxonomic-group distributions of positives (P) and negatives (N) in the Pintado-Grima benchmark and in the reported training datasets of phase-separation protein (PSP) predictors. Stacked bars show the fraction of proteins assigned to each major taxonomic group: Animals(M), Animals(NM), Fungi, Protists, Plants, Prokaryotes, and Viruses. Animals(M) denotes mammals, and Animals(NM) denotes non-mammalian animals. Distribution profiles were computed separately for positive and negative sets within each dataset. Predictors are grouped by the taxonomic scope of their training data (as indicated below): multi-taxon predictors (trained on multiple taxa) and single-taxon predictors (trained on a single taxon). **B and C** UMAP projection of ProtT5 sequence embeddings for proteins in the Pintado-Grima benchmark, colored by taxonomic group (B) and by class label (positive versus negative; C). **D and E** Performance of a logistic regression classifier trained to predict Pintado-Grima benchmark labels using only one-hot encoded taxonomic-group composition as input features. Receiver operating characteristic (ROC; D) and precision-recall (E) curves are shown. Shaded regions indicate variability across dataset splits generated with different random seeds, and insets report mean area under the curve (AUC) ± s.d. **F and G** Association between predictive performance and training-benchmark taxonomic composition distance. For each predictor, AUROC (F) and AUPRC (G) obtained directly from the Pintado-Grima benchmark are plotted against the Jensen-Shannon divergence between taxonomic composition of predictors’ training positive set and that of the benchmark (lower values indicate closer matching). Predictors were omitted when the Pintado-Grima benchmark did not report AUROC or AUPRC. Ellipses highlight the clustering of multi-taxon predictors (low divergence, higher performance; red ellipse) versus single-taxon predictors (higher divergence, reduced performance; blue ellipse).

To examine whether the taxonomic structure is reflected in sequence representation space, we calculated ProtT5^32^ embeddings of proteins in benchmark dataset and visualized them using UMAP^33^. Benchmark proteins segregated strongly by taxonomic group in the embedding space (Fig. 1B), indicating that taxon-linked sequence features dominate the low-dimensional geometry. Notably, the apparent separation between positives and negatives largely mirrored this taxon-driven organization (Fig. 1C), suggesting that the observed label separation can be largely explained by taxonomic structure.

If taxonomy indeed encodes label-associated information in this benchmark, then a classifier that uses only taxonomic identity should achieve non-trivial performance. We therefore trained a logistic regression model with one-hot encoded taxonomic-group indicators as the sole input features. Despite the complete absence of sequence-related features, this taxonomy-only baseline achieved measurable discrimination (mean AUROC = 0.856 ± 0.017; mean AUPRC = 0.579 ± 0.030 across random splits; Fig. 1D-E). These results provide direct evidence that taxonomic identity itself contains substantial label information in the Pintado-Grima benchmark, establishing taxonomy as a concrete confounder for predictor evaluation.

One practical consequence of this label-taxonomy coupling is that performance estimates may become sensitive to how closely a predictor’s training data match the benchmark’s taxonomic composition. Consistent with this, predictor performance on the Pintado-Grima benchmark was associated with training-benchmark taxonomic similarity: both AUROC and AUPRC tended to decrease as the Jensen-Shannon divergence^34,35^ between training and benchmark taxonomic distributions increased (Fig. 1F-G). Predictors could be broadly classified into two groups based on the taxonomic composition of their training datasets: multi-taxon predictors trained on multiple taxa, and single-taxon predictors trained on a single taxon (predominantly Animals(M)). Accordingly, multi-taxon predictors tended to occupy a low-divergence and high-performance regime, whereas single-taxon predictors more often showed higher divergence and reduced performance (Fig. 1F-G). Together with the taxon-structured embedding geometry (Fig. 1B-C) and the non-trivial performance of a taxonomy-only baseline (Fig. 1D-E), these results indicate that evaluation on the Pintado-Grima benchmark is intrinsically confounded by taxonomy, such that reported performance cannot be interpreted as a clean measure of PSP-discriminative ability.

Beyond taxonomic composition, predictors vary widely in training dataset size (Fig. S1A) and differ systematically in disorder composition (Fig. S1B-C), which can shape benchmark performance. In particular, the negative sets used to train multi-taxon predictors show IDR-proportion distributions that are closer to the Pintado-Grima benchmark negatives, whereas negatives from single-taxon predictors tend to exhibit higher IDR proportions (Fig. S1B). Likewise, the fraction of non-IDPs among multi-taxon negatives more closely matches the Pintado-Grima benchmark negative set (both exceeding 50%), while the corresponding positives remain overwhelmingly IDP-rich (non-IDP fraction <15%; Fig. S1C). By contrast, single-taxon predictors show a less pronounced disparity in IDP/non-IDP fraction between their positive and negative training sets (Fig. S1C). These patterns suggest that the apparent advantage of multi-taxon predictors on the Pintado-Grima benchmark may reflect not only taxonomic composition matching (Fig. 1F-G) but also closer matching of disorder composition, particularly within the negative set, to the benchmark. Together with the label-dependent taxonomic imbalance in the benchmark (Fig. 1A), these observations underscore the need for a taxonomy-aware benchmark that balances positives and negatives within each taxonomic group and matches the IDP/non-IDP ratio between labels, thereby forcing predictors to rely on LLPS-relevant features rather than taxonomy-or disorder-associated shortcuts.

### Construction of a taxonomy-aware, disorder-matched PSP benchmark dataset

Motivated by the strong coupling between PSP labels and taxonomic identity in the Pintado-Grima benchmark dataset and the IDP composition mismatches observed across training datasets, we constructed a new taxonomy-aware benchmark that explicitly controls two major confounders: (i) taxonomic composition and (ii) IDP/non-IDP composition (see Methods for details). Conceptually, the dataset was built such that (i) negatives are sampled to match the taxonomic-group composition of positives, and (ii) within each major taxonomic group, negatives are sampled to match the IDP/non-IDP composition of the corresponding positives (Table S1). An overview of the construction workflow is shown in Fig. 2 and summarized as a stepwise Sankey diagram in Fig. S2.

**Figure 2.**
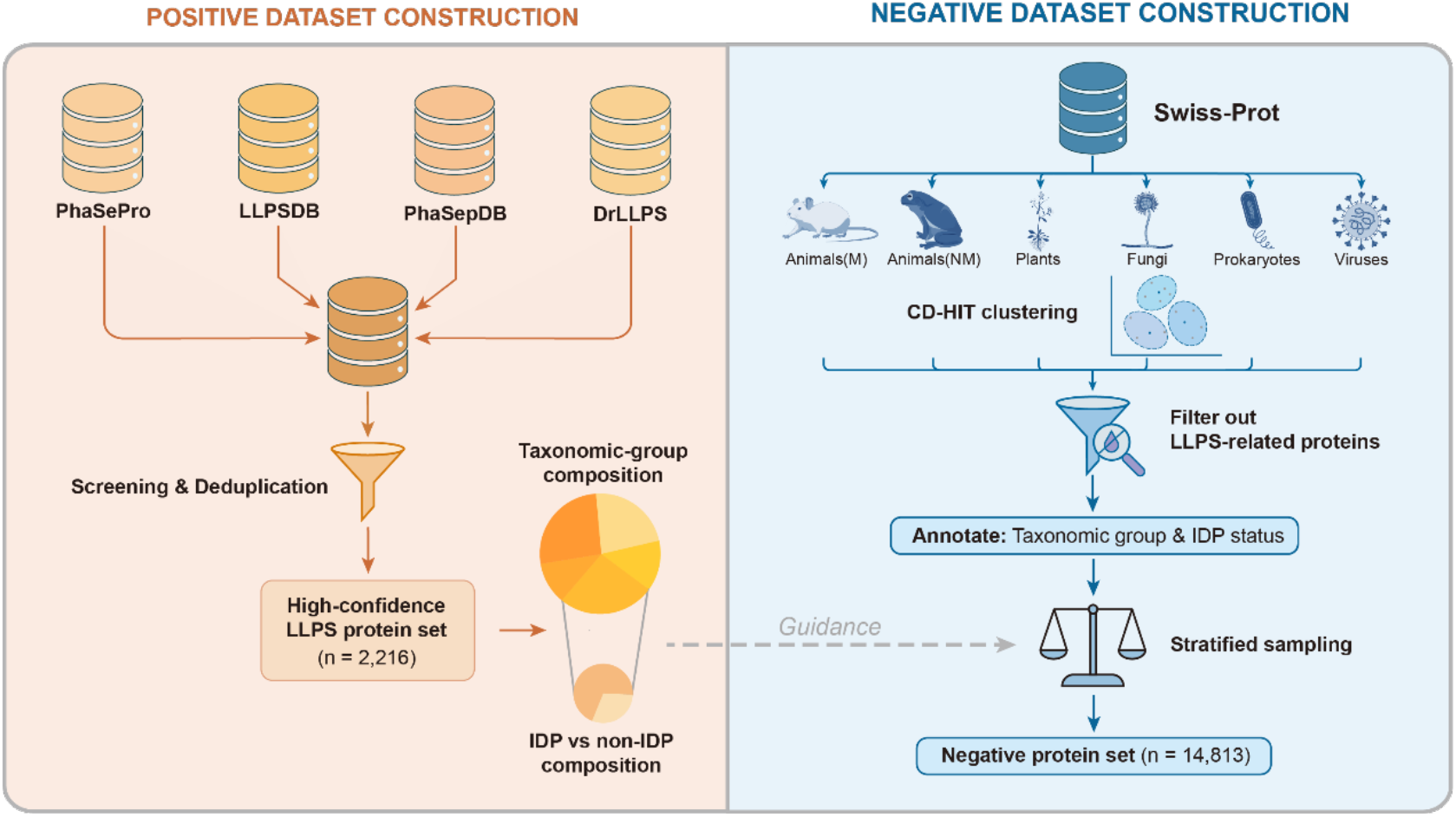
Taxonomy-aware construction of positive and negative datasets for PSP benchmarking. For the positive dataset, candidate proteins were compiled from four curated LLPS-focused resources (PhaSePro, LLPSDB, PhaSepDB and DrLLPS), merged, screened, and deduplicated to generate a high-confidence LLPS protein set (n = 2,216). The resulting taxonomic-group composition and the proportion of IDPs versus non-IDPs were summarized and used to guide negative-set sampling. For the negative dataset, candidate proteins were drawn from Swiss-Prot across major taxonomic groups (Animals(M), Animals(NM), Plants, Fungi, Prokaryotes and Viruses). Redundancy was reduced by CD-HIT clustering within each taxonomic group. Proteins with known or putative LLPS association were then removed, including the compiled PSPs and proteins showing sequence similarity to PSPs. The remaining proteins were annotated by taxonomic group and IDP status and selected by stratified sampling to match the positive set’s taxonomic-group and IDP/non-IDP composition, yielding the final negative protein set (n = 14,813). Representative species icons were obtained from BioGDP^38^.

We first assembled the positive set by merging entries from four curated LLPS-focused databases (LLPSDB^25^, PhaSePro^26^, PhaSepDB^27^, and DrLLPS^28^), followed by screening and de-duplication to obtain a high-confidence, non-redundant collection of 2,216 proteins spanning the major taxonomic groups (Animals(M), Animals(NM), Plants, Fungi, Prokaryotes, and Viruses; Fig. 2; Fig. S2A). Protists were excluded due to insufficient positives. To characterize how different resources contribute to the final positive compendium, we summarized cross-database overlap using an UpSet plot^36^ (Fig. S3A), which highlights both shared cores and complementary coverage across resources. Because our negative sampling is guided by the composition of the positive set, we explicitly quantified the positive set’s distribution across major taxonomic groups as well as its IDP versus non-IDP composition (Fig. 2; Fig. S3B). These summaries serve as the target composition that the negative set must match under taxonomy-aware sampling.

For the negative dataset, we compiled candidate proteins from Swiss-Prot across the same taxonomic groups as positives (Fig. 2; Fig. S2B). Within each taxonomic group, we first performed redundancy reduction via CD-HIT clustering^37^ to prevent over-representation of closely related sequences. To minimize contamination by true or putative PSPs (and to reduce information leakage through close homologs), we then removed proteins with known or suspected LLPS association. This filtering step included (i) the compiled PSP positives themselves and (ii) additional proteins exhibiting sequence similarity to PSPs. The resulting filtered pool was annotated by taxonomic group and IDP status, enabling stratified sampling to match the positive set’s composition. This procedure yielded a final negative set of 14,813 proteins. By design, this taxonomy-aware, disorder-matched benchmark minimizes taxonomy- and disorder-based shortcuts, making cross-taxon comparisons and taxonomy-stratified PSP benchmarking more interpretable.

### Taxon-specific baselines shape LLPS-associated sequence and biophysical features

Using our taxonomy-aware, disorder-matched benchmark, we next examined how sequence and biophysical features associated with PSPs vary across taxa. To this end, we analyzed the PSPs in the positive set in a taxonomy-stratified framework. We first examined whether PSP sequence composition is conserved across taxa. For each amino acid, we computed the relative change in mean residue frequency within a given taxon compared with the global mean across all taxa (Fig. 3A). The resulting heatmap is far from uniform: each taxon exhibits a distinct signed pattern across residues, and many amino acids flip direction (enriched in one taxon but depleted in another). For example, Prokaryotes PSPs show a striking enrichment of aliphatic residues, with alanine, valine, leucine, and isoleucine all strongly enriched, whereas Fungi PSPs display a split aliphatic signature in which isoleucine is strongly enriched but alanine and valine are clearly depleted. Even within animals, Animals(NM) PSPs show striking depletion of tryptophan and enrichment of glycine, while Animals(M) PSPs display comparatively modest deviations. To quantify overall compositional divergence from the mammalian reference, we computed the Jensen-Shannon divergence (JSD) between each taxon’s amino-acid composition profile and that of Animals(M), which revealed appreciable divergence across taxa (Fig. 3B).

**Figure 3.**
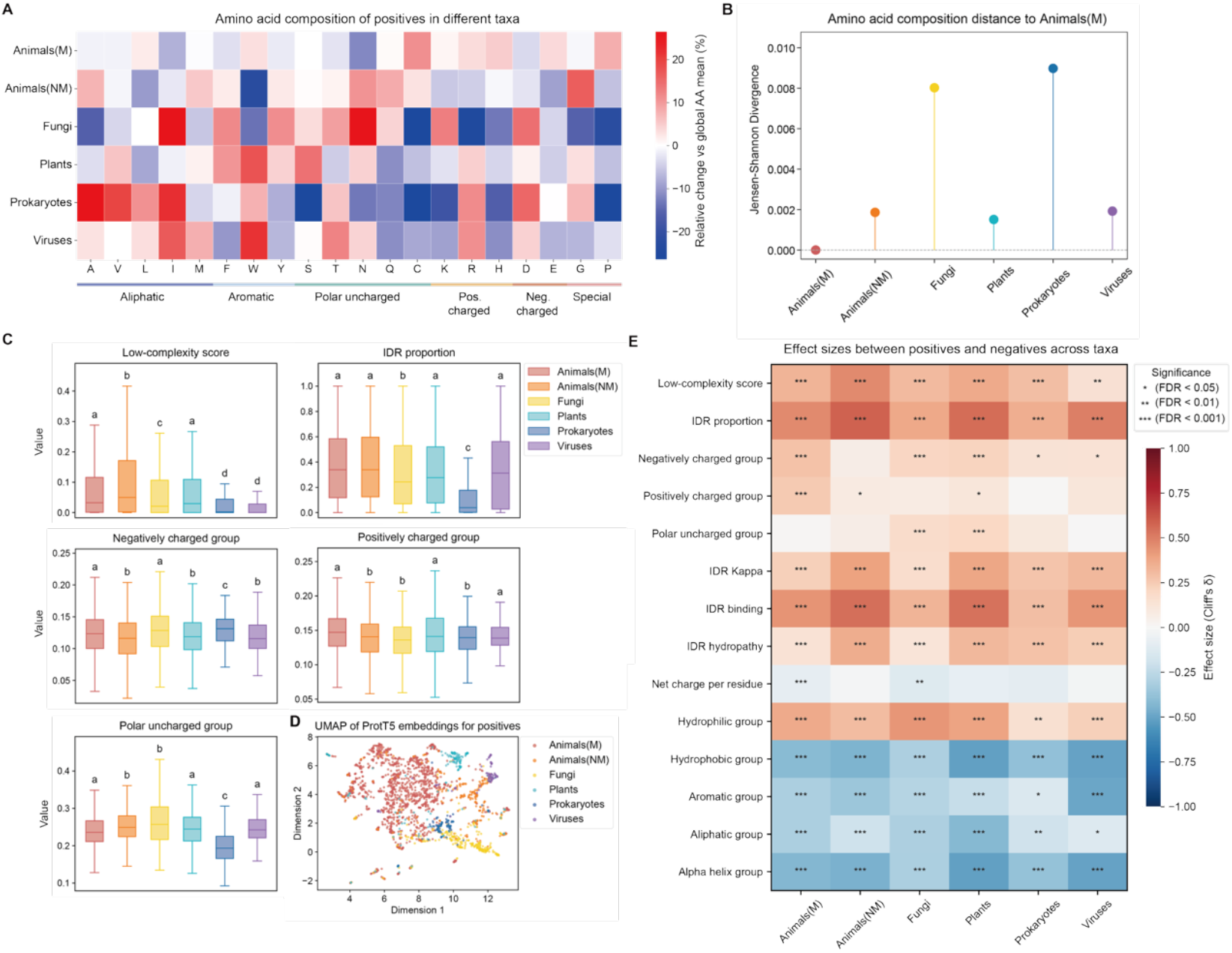
Taxon-dependent sequence and biophysical properties of PSPs reflect distinct proteome baselines. **A** Amino-acid composition of PSPs across taxonomic groups. Heatmap entries represent the relative change (%) of each amino acid’s mean frequency within a taxonomic group compared with the global mean across all proteins. Red indicates enrichment and blue indicates depletion. Amino acids are grouped by physicochemical class (aliphatic, aromatic, polar uncharged, positively charged, negatively charged and special). **B** Taxonomic distance in amino-acid composition relative to Animals(M). Points show the Jensen-Shannon divergence between each taxon’s amino-acid composition profile and that of Animals(M); the dashed line denotes zero divergence. **C** Distributions of representative sequence/biophysical features for positives across taxa, including low-complexity score, intrinsically disordered region (IDR) proportion, the fraction of negatively charged residues (D, E), the fraction of positively charged residues (K, R, H) and the fraction of polar uncharged residues (S, T, N, Q, C). Boxplots show median (central line), interquartile range (IQR; 25th-75th percentiles), and whiskers extending to the most extreme values within 1.5 × IQR. Letters denote compact letter display (CLD) groupings derived from two-sided Mann-Whitney U tests with Benjamini-Hochberg false discovery rate (BH-FDR) correction; groups sharing at least one letter are not significantly different, whereas groups with no letters in common differ significantly (BH-FDR < 0.05). **D** UMAP projection of ProtT5 embeddings for positive proteins, colored by taxonomic group. **E** Effect sizes between PSPs and proteome background across taxa. Heatmap shows the taxon-stratified effect size (Cliff’s δ) for representative sequence/biophysical features. Color intensity reflects the magnitude of the effect size. Asterisks denote statistical significance from two-sided Mann-Whitney U tests with Benjamini-Hochberg false discovery rate (BH-FDR) correction; the number of asterisks indicates the FDR tier (*** indicates FDR < 0.001, ** indicates FDR < 0.01, and * indicates FDR < 0.05).

We then examined whether commonly used LLPS-associated properties show similar taxon dependence. We quantified a panel of disorder-, charge-, and composition-related descriptors, with Fig. 3C showing representative features (low-complexity score, IDR proportion, and the fractions of negatively charged, positively charged, and polar uncharged residues), and Fig. S4 extending the analysis to additional IDR-centric and compositional properties (IDR κ^39^, IDR binding score^40,41^, IDR hydropathy^39^, NCPR, and multiple residue-group fractions) (see Methods for details). The analysis revealed that feature distributions were broadly taxon dependent (Fig. 3C; Fig. S4). A prominent example is the contrast between Prokaryotes and Animals(M): relative to Animals(M), Prokaryotes PSPs exhibit markedly lower IDR proportion and low-complexity scores, reduced IDR κ and IDR binding scores, together with strikingly higher hydrophobic and aliphatic fractions and an elevated α-helix-associated residue fraction. Consistent with these feature-level heterogeneity, a UMAP projection^33^ of ProtT5 embeddings^32^ of PSPs revealed clear taxon-linked organization (Fig. 3D).

Since many sequence properties vary across taxa at the proteome level, we further examined whether the heterogeneity observed above could be explained simply by taxon-specific proteome baselines. We therefore performed a within-taxon background-contrast analysis: for each taxonomic group, we compared PSPs against the corresponding taxon-specific proteome background and quantified PSP-background separation for each feature using Cliff’s δ^42^ (Fig. 3E). While some feature-taxon pairs show weak or non-significant separation, for a majority of features, PSPs deviate from their own proteome baselines in a consistent direction (Fig. 3E). This indicates that much of the apparent cross-taxon heterogeneity in absolute feature values reflects taxon-specific proteome baselines, whereas LLPS-associated feature shifts relative to background are comparatively conserved, highlighting that absolute feature levels are not directly comparable across taxa.

We next characterized the functional roles and domain architectures of PSPs in each taxonomic group. We performed Gene Ontology (GO) and protein domain enrichment analyses for PSPs from each taxon. Both GO and Pfam enrichment landscapes were strongly taxon specific, with distinct domain families and functional themes enriched in different groups (Fig. S5A-B). For instance, Animal(M) PSPs were enriched for RNA-associated domains such as RRM, KH, and DEAD/DEAH-box helicase modules, whereas Prokaryotes showed strong enrichment for Rubisco-related domains (Fig. S5A). GO enrichments mirrored these domain-level shifts (Fig. S5B). Together with the proteome-contrast analysis, these results indicate that taxon-specific proteome baselines, functional niches, and domain architectures shape how PSPs implement LLPS in different lineages, reinforcing the need to evaluate PSP predictors in a taxon-resolved rather than pooled multi-taxon setting.

### Taxonomic- and disorder-dependent performance of PSP predictors

Using our taxonomy-aware, disorder-matched benchmark, we compared PSP predictors in six taxonomic groups: Animals(M), Animals(NM), Plants, Fungi, Prokaryotes, and Viruses. Twenty predictors available before January 2026 (see Methods for details) were evaluated and grouped into two main categories (Table S2): (i) rule-based, unsupervised models (PLAAC^5^, PScore^7^, and PSPer^8^); and (ii) machine-learning (ML)-based supervised models. The supervised models were further subdivided by input representation into engineered-feature models (CatGRANULE^6^, FuzDrop^9^, PSAP^12^, PhaSePred^13^, PICNIC^15^, PSPire^16^, FLFB^17^, and ROBOT^24^), protein language model (PLM) embedding-based models (Seq2Phase^18^, PSTP^20^, PhaseNet^21^, MambaPhase^22^, and PSPsPredict^23^), and non-PLM representation-learning models (Droppler^10^, DeePhase^11^, PSPredictor^14^, and PSPHunter^19^). Predictor performance across taxonomic groups was quantified using both threshold-free metrics (AUROC and AUPRC) and threshold-dependent metrics (MCC, F1-score, precision, recall, specificity, and accuracy) (see Methods for details; Table S3).

Across these six taxonomic groups, both AUROC- and AUPRC-based rankings varied markedly (Fig. 4A). Unless otherwise stated, we refer to AUROC-based ranking in the text, but AUPRC showed broadly similar patterns. Rule-based, unsupervised predictors generally underperformed relative to ML-based supervised models, suggesting that fixed heuristics capture only a limited subset of LLPS-relevant signals. Among supervised predictors, PLM embedding-based models (Seq2Phase, PSTP, PhaseNet, MambaPhase and PSPsPredict) typically achieved the strongest performance, frequently ranking among the top five across multiple taxa, indicating that PLM-derived embeddings provide rich sequence and biophysical representations that benefit PSP prediction. Several engineered-feature models (PSPire, ROBOT, and PICNIC) performed well in one or two taxa, while the non-PLM representation-learning models DeePhase and PSPHunter each ranked in the top five in two groups. These taxon-resolved results underscore that a single aggregate score on a pooled multi-taxon dataset (pooled evaluation, see Fig. S6A) can obscure substantial differences in behavior across taxa and may be dominated by mixture effects rather than stable generalization across taxonomic groups. Threshold-dependent metrics showed similar patterns (Table S3).

**Figure 4.**
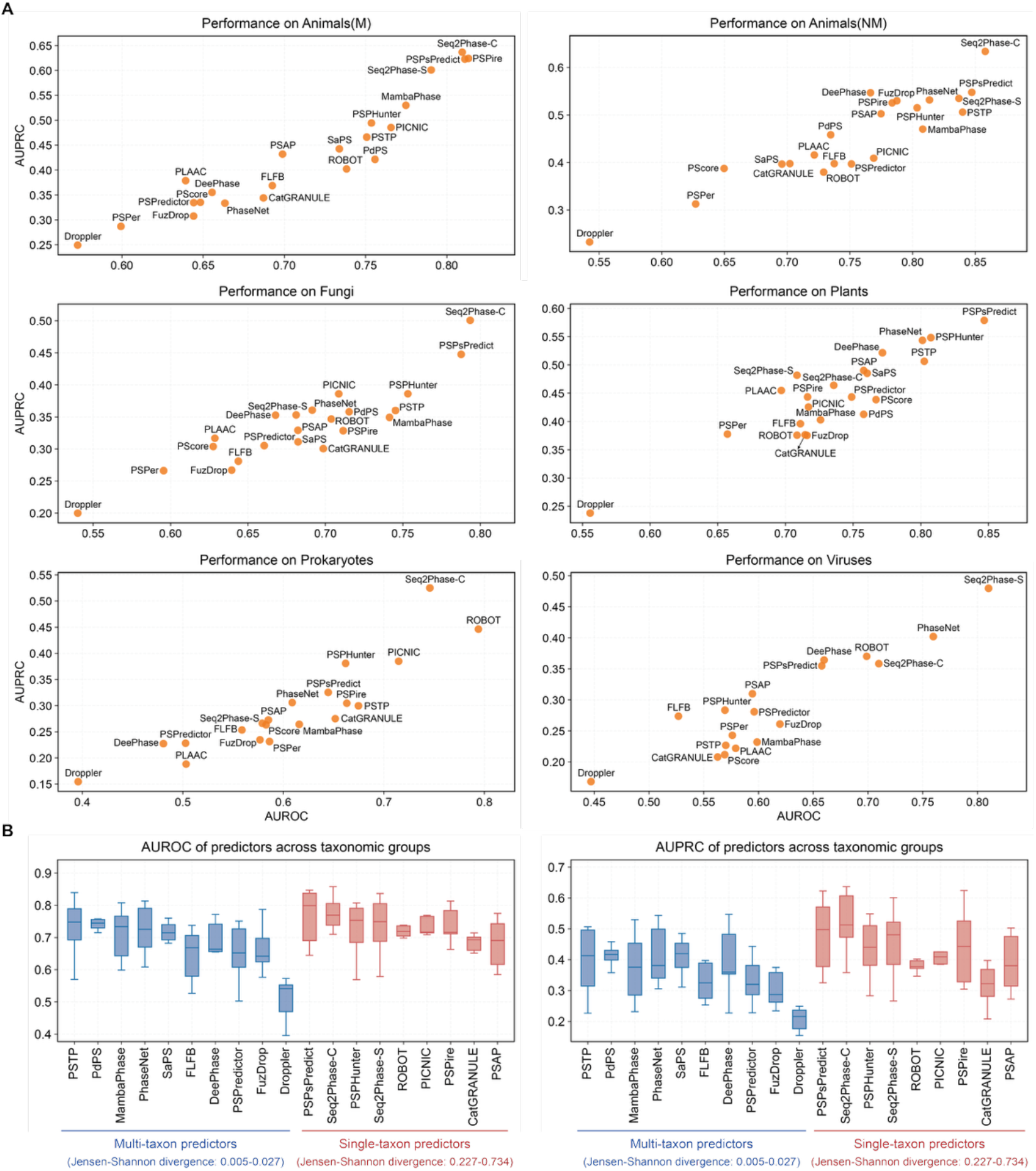
Performance of PSP predictors on the benchmark dataset. **A** Scatter plots show AUROC versus AUPRC, with each point labelled by predictor name. The two PhaSePred variants (SaPS for self-assembling proteins and PdPS for partner-dependent proteins) and the two Seq2Phase variants (Seq2Phase-C targeting client proteins and Seq2Phase-S targeting scaffold proteins) were evaluated separately. Performance was evaluated separately across six taxonomic groups: Animals(M), Animals(NM), Plants, Fungi, Prokaryotes, and Viruses. The two PhaSePred variants (SaPS and PdPS) was excluded from evalution on Prokaryotes subset due to limited protein support; while PhaSePred, PSPire and PICNIC were excluded from evalution on Viruses subset for the same reason. **B** Distribution of AUROC and AUPRC of predictors across taxonomic groups. Predictors are grouped by the taxonomic scope of their training data (as indicated below): multi-taxon predictors (trained on multiple taxa) and single-taxon predictors (trained on a single taxon). Since the three rule-based, unsupervised models (PLAAC, PScore, and PSPer) do not require training data, they were not shown. Jensen-Shannon divergence was calculated between taxonomic composition of predictors’ training positive set and that of the benchmark (lower values indicate closer matching). Boxplots show median (central line), interquartile range (IQR; 25th-75th percentiles), and whiskers extending to the most extreme values within 1.5 × IQR.

On our taxonomy-aware, disorder-matched benchmark, the dependence of predictor performance on training-benchmark taxonomic similarity observed in the Pintado-Grima benchmark (Fig. 1F-G) was absent. Multi-taxon and single-taxon predictors exhibited broadly overlapping AUROC and AUPRC distributions across taxonomic groups, and we did not observe systematic relationship between performance and the Jensen-Shannon divergence between training and benchmark taxonomic distributions (Fig. 4B). These results indicated that evaluation based on our benchmark dataset is no longer driven by simple taxonomic matching between training data and benchmark composition.

Our previous study showed that PSP predictors can behave differently on PSPs with IDRs (ID-PSPs) versus those without IDRs (noID-PSPs)^16^. Therefore, we stratified positives by disorder status to quantify performance gaps between these two classes. We evaluated ID-PSP and noID-PSP performance in Animals(M), as the remaining taxonomic groups contained too few noID-PSPs to support reliable estimation. Across all predictors tested, identifying noID-PSPs was consistently more challenging than identifying ID-PSPs, yielding a uniform decrease in both AUROC and AUPRC (Fig. S6B-C). The magnitude of this performance gap varied substantially across predictors, indicating that current predictors differ widely in their ability to handle noID-PSPs. Seq2Phase and PSPire achieved the highest performance on noID-PSPs, whereas several predictors showed limited discrimination in this setting. Together, these results indicate that noID-PSPs represent a distinct and more challenging regime for current predictors, motivating routine disorder-stratified reporting for fair interpretation.

## Discussion

In computational biology, benchmarks are often treated as neutral arbiters of method quality. The recently released Pintado-Grima benchmark^29^ has begun to play this role for PSP predictors. However, quantitative analysis of this dataset reveals systematic departures from such neutrality. We find that this mixed-taxon benchmark couples PSP labels to taxonomic identity and disorder composition, and that models can achieve high scores by exploiting benchmark-specific shortcuts rather than capturing transferable LLPS-associated grammars. Consistent with this, the Pintado-Grima benchmark can systematically favour predictors trained on multi-taxon datasets over single-taxon predictors, independent of their performance within their intended taxonomic group. To address this, our taxonomy-aware, disorder-matched benchmark is designed explicitly to minimize these effects, and reveals substantial taxon- and disorder-dependent variation in PSP predictor performance that pooled, shortcut-prone evaluations obscure.

In practice, studies that introduce a new predictor typically enforce that their independent test set is non-redundant with respect to their own training data, often considering both positive and negative proteins. However, when a benchmark aims to evaluate multiple previously published predictors, it is rarely feasible to guarantee simultaneous exclusion of all training sets. Training lists are frequently incomplete, inconsistently reported, or unavailable, making comprehensive decontamination impractical. In this work, to improve fairness while maintaining a realistic evaluation setting, we curated and removed the training positive sets of all predictors for which training data were obtainable; for methods where training positives were not publicly released, we attempted to obtain them by contacting the authors (see Methods for details). In contrast, we did not remove training negatives, because for some predictors the negative set is extremely large and effectively defined as the remainder of the proteome (for example, PSAP^12^ treats the non-positive human proteome as negatives, totaling ∼19,000 proteins). Removing such negatives would eliminate a substantial fraction of the candidate negative pool and, more importantly, would alter its taxonomic and disorder composition, confounding the very benchmark conditions we aim to control. We therefore prioritized positive-set decontamination as the most direct safeguard against optimistic inflation, while acknowledging that some overlap in negative training proteins remains an inherent limitation of multi-method benchmarking.

Our benchmark results also suggest practical directions for future PSP predictor design. The consistent underperformance of rule-based approaches implies that fixed heuristic rules capture only a limited subset of relevant biophysical signals. By contrast, predictors based on PLM embeddings exhibit comparatively robust performance across taxonomic groups, supporting the use of PLM-derived representations as a strong default backbone for PSP prediction. At the same time, the good performance of PSPire^16^ on mammals highlights the added value of context-dependent regulatory features, such as post-translational modification annotations. Together, these observations motivate a hybrid architecture in which PLM embeddings provide the core representation for cross-taxon generalization, while lightweight modules incorporate regulatory and cellular-context features where appropriate.

## Methods

### Taxonomic composition analysis

To characterize taxonomic composition, we analyzed the benchmark dataset introduced by Pintado-Grima et al.^29^ (2,876 proteins), together with the training datasets of predictors evaluated in that study. Training datasets were collected from original publications, supplementary materials, or publicly available repositories when available. As training datasets of Droppler^10^ and FLFB^17^ could not be obtained, we contacted the authors and received the FLFB training data which were provided as sequence fragments, but did not receive a response from the Droppler authors. Accordingly, Droppler was excluded from the taxonomic composition profiling. For FLFB, sequence fragments were mapped to full-length proteins by BLASTP against Swiss-Prot. Fragments that were fully contained within a single protein and produced a unique match were assigned to that protein and used for downstream analyses. In addition, the four rule-based and unsupervised models (PLAAC^5^, PScore^7^, PSPer^8^, and R+Y^43^) were excluded because no explicit training datasets are defined for these methods. For PSPredictor^14^, whose training data were also provided as sequence fragments, we mapped fragments to full-length proteins using the same BLASTP-based procedure applied to FLFB.

Taxonomic annotations were retrieved from UniProtKB and mapped to higher-level taxonomic lineages. Each protein was assigned to one of seven taxonomic groups: mammals (Animals(M)), non-mammalian animals (Animals(NM)), plants, fungi, protists, prokaryotes, or viruses. Taxonomic distribution profiles were computed separately for the positive and negative sets within each dataset.

### Taxon-only classification model on the Pintado-Grima benchmark dataset

To quantify the predictive power of taxonomic information alone on the Pintado-Grima benchmark, we trained a classifier using only taxonomic group labels as input. Each protein was encoded as a one-hot vector indicating its assigned taxonomic group, and a logistic regression model was fit to predict class labels. Performance was evaluated by randomly splitting the benchmark dataset into training and test sets at an 8:2 ratio. This procedure was repeated for ten independent splits using different random seeds. Predictive performance was summarized using the area under the receiver operating characteristic curve (AUROC) and the area under the precision-recall curve (AUPRC).

### Intrinsically disordered regions calculation

Intrinsically disordered regions (IDRs) were generated by first computing residue-level disorder propensity scores with AIUPred^44,45^ and then applying a post-processing procedure following the MobiDB-lite protocol^46^. AIUPred scores were binarized using a disorder threshold of 0.5, yielding an initial residue-wise disorder state. To reduce spurious fragmentation of disordered segments, morphological smoothing was applied using consecutive dilation and erosion operations, followed by gap-merging to connect long disordered regions separated by short structured intervals. Long IDRs were then defined as contiguous disordered segments with a minimum length of 20 residues after post-processing. Proteins harboring at least one long IDR were classified as intrinsically disordered proteins (IDPs), whereas proteins lacking any long IDR were classified as non-IDPs. Likewise, PSPs with at least one long IDR were designated ID-PSPs, while PSPs without long IDRs were designated noID-PSPs.

### Positive dataset construction

Phase separating positive proteins were collected by integrating four curated LLPS-focused databases, including LLPSDB v2.0^25^, PhaSePro v1.1.0^26^, PhaSepDB v3.0^27^, and DrLLPS^28^. CD-CODE^47^ was not used as a primary source to compile phase-separating proteins because it is a condensate-centric, community-editable “living” knowledgebase in which entries are continuously updated by contributors and maintainers, and its protein annotations primarily reflect condensate association rather than an intrinsic or autonomous phase-separation capability. Proteins shorter than 50 amino acids or longer than 3,000 amino acids were excluded. Proteins containing uncommon amino acids such as “BJOUXZ” were also excluded.

The combined positive records were subjected to systematic screening and deduplication. For LLPSDB, proteins from “Ambiguous” subset were excluded. For DrLLPS, only proteins with condensate information and tissue/cell annotations were included. Proteins reported in publications covering more than 10 entries from DrLLPS were excluded to avoid possible high-throughput annotations. Redundant records across databases were then collapsed to a non-redundant protein set, and the resulting taxonomic composition of the set was quantified. Protists were excluded from the set due to insufficient number of high-confidence phase separating positive proteins. A final set of 2,216 high-confidence PSPs was obtained and used as the positive benchmark set (Fig. 2; Fig. S2A).

### Negative dataset construction

Candidate negative proteins were derived from the Swiss-Prot database with sequence length from 50 to 3000 amino acids. To minimize contamination by proteins with reported condensate association, candidates showing sequence similarity to any protein included in the LLPS-focused databases used for positive set construction, as well as CD-CODE v1^47^, were removed based on BLASTP alignments using thresholds of sequence identity ≥40% and alignment coverage ≥70%. CD-CODE was used as an additional exclusion resource to reduce the risk that proteins with any reported condensate association were retained as negatives, thereby enforcing a stringent definition of the negative benchmark set. Candidates containing uncommon amino acids such as “BJOUXZ” were further excluded. The remaining proteins were classified into taxonomic groups. To reduce sequence redundancy while preserving taxon-specific diversity, CD-HIT clustering was performed independently within each taxonomic group using a sequence identity cutoff of 0.4.

Final negative set construction was performed by stratified sampling, with target counts guided by the taxonomic group and IDP/non-IDP composition of the positive set. Within each stratum, specific proteins were selected using an embedding-based diversity sampling strategy. Mean-pooled ESM-2 embeddings^48^ were used to represent protein sequences, and a cosine-distance-based k-center greedy algorithm was applied to select proteins that maximally covered the sequence embedding space. This procedure ensured that the negative dataset spanned a broad and diverse sequence space while maintaining matched taxonomic group and disorder profiles relative to the positive set. The final negative benchmark set consisted of 14,813 proteins (Fig. 2; Fig. S2B).

### Taxon-stratified characterization of LLPS-associated features

The LLPS-associated features were calculated for proteins in the constructed positive set. Firstly, amino-acid composition profiles were computed separately for each taxonomic group by calculating the mean residue frequency of each amino acid across proteins in that group. To highlight taxon-specific deviations, relative changes were computed with the following formula:

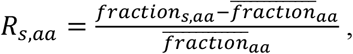

where *fraction*_*s,aa*_ represents the mean residue frequency of amino acid (aa) in taxon s, 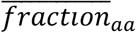 means the average aa frequency of all protein samples.

In addition to amino-acid composition, fractions of the following amino acid groups were calculated: Aliphatic (A, V, L, I, M), Aromatic (F, W, Y), Polar uncharged (S, T, N, Q, C), Positively charged (K, R, H), Negatively charged (D, E), Hydrophilic (S, T, H, N, Q, E, D, K, R), Hydrophobic (V, I, L, F, W, Y, M), and Alpha Helix (V, I, Y, F, W, L). The fractions were defined as residue counts normalized by protein length.

A panel of disorder- and charge-related features were further calculated. Intrinsically disordered region (IDR) proportion was defined as the total length of all IDRs divided by protein length. The IDR binding score was defined as the proportion of residues whose ANCHOR^40^ score exceeded 0.5 within IDRs, normalized by the full sequence length. IDR hydropathy was computed using the normalized Kyte-Doolittle hydrophobicity scale^49^ implemented in localCIDER^39^. Net charge per residue (NCPR) was defined as the sequence-length-normalized net charge, computed by subtracting the counts of negatively charged residues (D and E) from positively charged residues (K, R, and H). The IDR Kappa (κ; charge patterning) was defined as the maximum κ value computed by localCIDER^39^ across all IDRs identified in each protein.

Additionally, low-complexity score was calculated following the protocol used by PSAP^12^. Briefly, for each residue, a sliding window of 20 amino acids was constructed with the residue positioned at the window center. The number of unique amino-acid types within the window was then counted, and the central residue was assigned a low-complexity label of 1 when the unique count was ≤7, and 0 otherwise. For each protein, the low-complexity score was defined as the total number of low-complexity-labeled residues normalized by protein length.

To distinguish LLPS-associated signatures from taxon-specific proteome baselines, PSPs were compared against taxon-matched background proteins used for CD-HIT clustering, comprising 446,887 proteins in total (Fig. S2B). Feature separation within each taxonomic group was quantified using Cliff’s δ, a rank-based effect size: δ > 0 indicates that PSPs tend to take higher values than the background, whereas δ < 0 indicates a shift toward lower values. Statistical significance was assessed using two-sided Mann-Whitney U tests with Benjamini-Hochberg false discovery rate (BH-FDR) correction.

### Taxonomy-aware benchmarking of PSP predictors

PSP predictors available before January 2026 were collected for benchmarking. A total of twenty predictors including PLAAC^5^, CatGRANULE^6^, PScore^7^, PSPer^8^, FuzDrop^9^, Droppler^10^, DeePhase^11^, PSAP^12^, PhaSePred^13^, PSPredictor^14^, PICNIC^15^, PSPire^16^, FLFB^17^, Seq2Phase^18^, PSPHunter^19^, PSTP^20^, PhaseNet^21^, MambaPhase^22^, PSPsPredict^23^, and ROBOT^24^ were evaluated. The two PhaSePred variants (SaPS for self-assembling proteins and PdPS for partner-dependent proteins) and the two Seq2Phase variants (Seq2Phase-C targeting client proteins and Seq2Phase-S targeting scaffold proteins) were evaluated separately. Since R+Y^43^ is a rule-based method that predicts saturation concentration (csat) for FUS-family-like disordered proteins from the abundance of arginine (R) and tyrosine (Y) residues, it was excluded from evaluation. For predictors providing executable models or programmatic interfaces, predictions were generated according to published instructions. For predictors available only via web servers, prediction results were retrieved using automated Selenium-based scripts. For predictors providing only precomputed proteome-wide predictions without an executable pipeline, the published results were used directly. For PSTP, we used the mixed-dataset-trained model and took its sequence-level prediction score as the final phase-separation propensity for each protein. For PSPire, we used the version including the phosphorylation frequency feature for human proteins and the version excluding this feature for non-human species.

Benchmark evaluation was performed on the six taxonomic groups of benchmark dataset: Animals(M), Animals(NM), Plants, Fungi, Prokaryotes, and Viruses. The two PhaSePred variants (SaPS and PdPS) was excluded from evalution on Prokaryotes subset due to limited protein support; while PhaSePred, PSPire and PICNIC were excluded from evalution on Viruses subset for the same reason. In addition, evaluation was performed on a pooled benchmark subset combining Animals(M), Animals(NM), Plants, and Fungi. Prokaryotes and Viruses subsets were excluded from pooled analyses due to limited predictor support.

To ensure fair comparison, proteins not supported by any predictor were excluded from the constructed evaluation sets. To mitigate potential training-data leakage and further improve benchmarking fairness, proteins appearing in the available training positive sets of evaluated predictors were removed from the evaluation sets. Because no response was received from the Droppler authors, no explicit training-positive exclusion could be applied for Droppler. After excluding training-positive proteins, class balance varied substantially across taxonomic groups. To make performance comparable across groups, we set the target class balance to the minimum negative-to-positive ratio observed among all taxonomic groups, and randomly downsampled negatives in the other groups to match this ratio. This procedure was repeated ten times using different random seeds. In each repeat, the sampled negatives were additionally constrained to match the IDP/non-IDP composition of the positive set within the same taxonomic group. Performance metrics were averaged across the ten repeats.

Predictor performance across taxonomic groups was quantified using both threshold-free metrics (AUROC and AUPRC) and threshold-dependent metrics (MCC, F1-score, precision, recall, specificity, and accuracy). Calculation of threshold-dependent metrics requires binarization of continuous predictor outputs. When explicit decision thresholds were defined by the original authors, those thresholds were applied: 0.75 for CatGRANULE, 4 for PScore, and 0.6 for FuzDrop. For PLAAC, proteins with a COREscore (COREscore not NaN) were classified as positive. For FLFB, proteins assigned the output label “LLPS” were classified as positive. For the remaining predictors (PSPer, Droppler, DeePhase, PSAP, PhaSePred, PSPredictor, PICNIC, PSPire, Seq2Phase, PSPHunter, PSTP, PhaseNet, MambaPhase, PSPsPredict, and ROBOT), for which explicit thresholds were not provided, the decision threshold that maximized MCC on the corresponding evaluation set was used. The threshold yielding the maximum MCC was obtained by scanning thresholds from 0.01 to 0.99 in steps of 0.01. Thresholds were determined independently for each predictor across ten independently generated negative-set subsamples.

Additionally, within each taxonomic group, positives were further stratified by disorder status into ID-PSPs and noID-PSPs to quantify performance differences between these two classes. Performance on ID-PSPs and noID-PSPs was evaluated only in the Animals(M) subset, because the other subsets contained too few noID-PSPs (fewer than 30) to support reliable evaluation.

## Data availability

All datasets used in this study are publicly available and detailed in Supplementary Data. The following four databases were used for the collection of phase-separating protein datasets: LLPSDB v2.0^25^, PhaSePro v1.1.0^26^, PhaSepDB v3.0^27^, and DrLLPS^28^.

## Acknowledgements

This work was supported by the National Natural Science Foundation of China (32325012, 32488101), the National Key Research and Development Program of China (2021YFA1302500), the Science and Technology Commission of Shanghai Municipality (23JS1401200, 24YF2734900), and the China Postdoctoral Science Foundation (2024M762403).

## Author contributions

Y.Z. conceived the project; S.H. and H.S. conducted data analysis, benchmark dataset construction and performance evaluation; Y.Z., S.H., and H.S. wrote the manuscript.

