## Supplementary figures and tables for "Taxonomy-aware, disorder-matched benchmarking of phase-separating protein predictors"

[#]These authors contributed equally

### SUPPLEMENTARY FIGURES

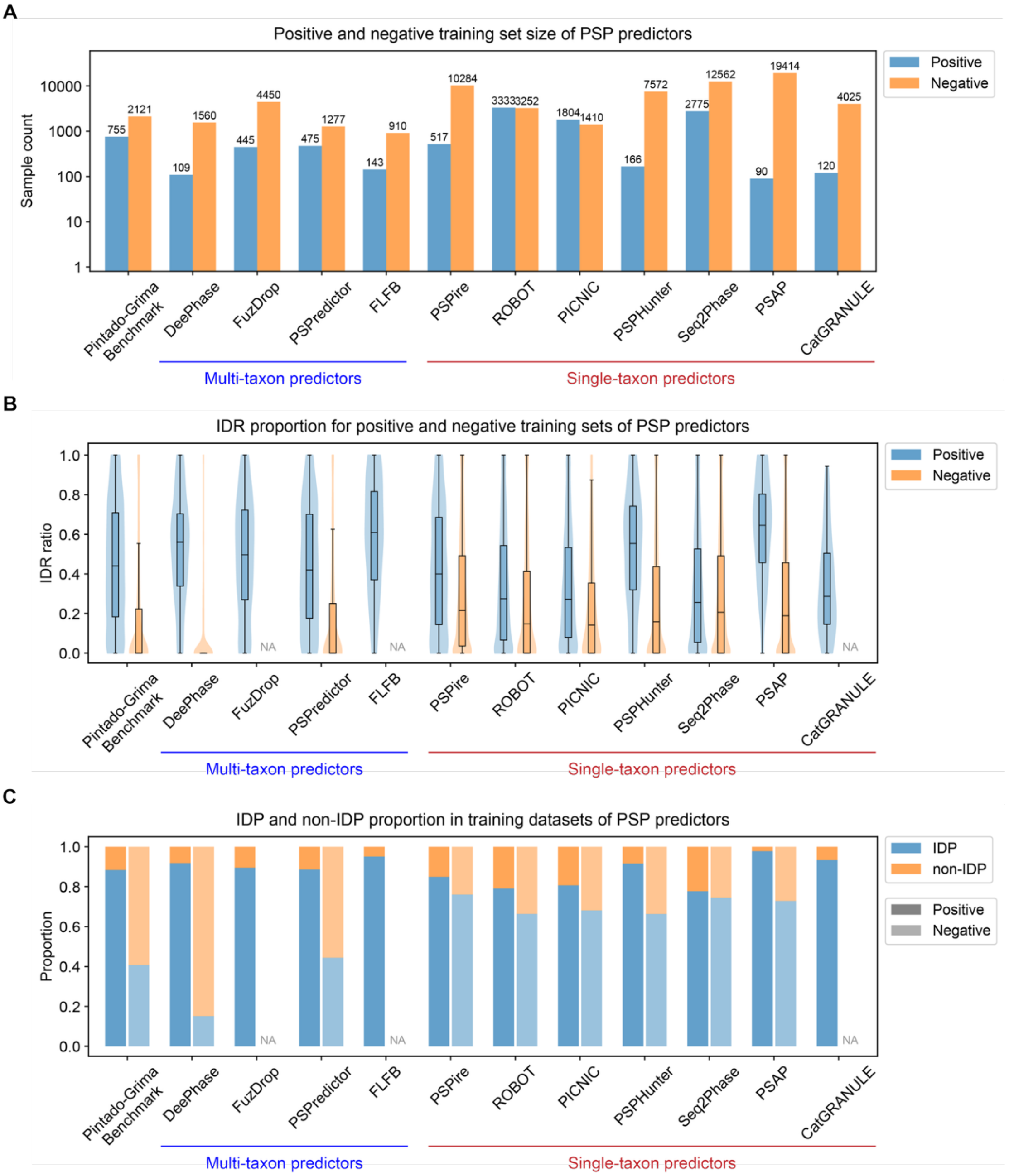


**Supplementary Figure 1**. Training-set size, intrinsically disordered regions (IDRs) content, and proportion of proteins with IDRs (IDPs) vary widely across PSP predictors and the Pintado-Grima benchmark. Predictors are grouped by the taxonomic scope of their training data (as indicated below): multi-taxon predictors (trained on multiple taxa) and single-taxon predictors (trained on a single taxon). **A** Sample sizes of the positive and negative training sets for PSP predictors and the Pintado-Grima benchmark. Bar heights indicate sample counts (log-scaled y axis), with counts annotated above bars. **B** Distributions of IDR proportion for proteins in the positive and negative training sets for PSP predictors and the Pintado-Grima benchmark. Violin plots show the full distribution with an overlaid boxplot. The central line indicates the median. The box represents the interquartile range (IQR), covering the 25th to 75th percentile. Whiskers extend to the data's minima and maxima within 1.5 × IQR. "NA" indicates that the corresponding set was not available for analysis. **C** Proportions of IDP and non-IDP for proteins in the positive and negative training sets for PSP predictors and the Pintado-Grima benchmark. Stacked bars show the fraction of proteins classified as IDP versus non-IDP within each set. "NA" indicates that the corresponding set was not available for analysis.

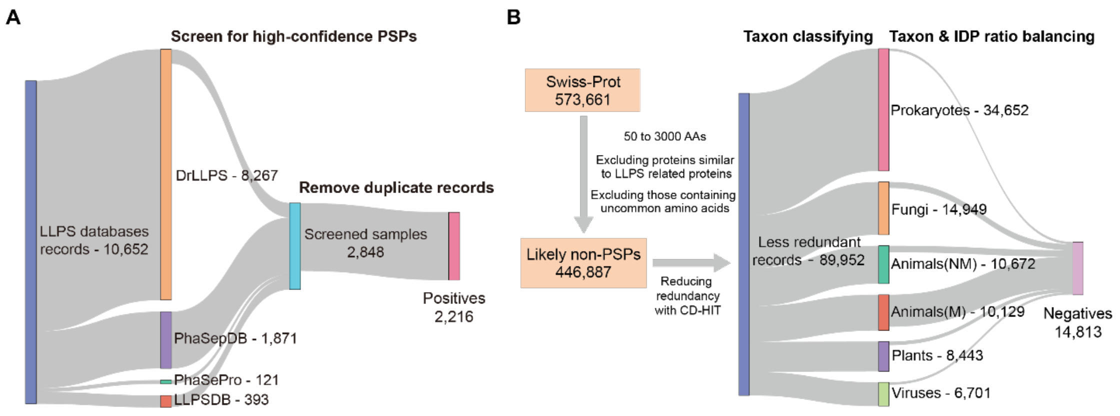


**Supplementary Figure 2**. Overview of taxonomy-aware benchmark dataset construction. Sankey diagram summarizing the workflow used to derive high-confidence positives and a taxon-matched negative pool. **A** For the positive dataset, records from curated LLPS-focused resources (DrLLPS, PhaSepDB, PhaSePro and LLPSDB) were merged, screened and deduplicated to obtain the final positive set (n = 2,216). **B** For the negative dataset, Swiss-Prot proteins were filtered to obtain a pool of likely non-PSPs, followed by redundancy reduction within each taxonomic group (Animals(M), Animals(NM), Plants, Fungi, Prokaryotes and Viruses) to enable taxonomy-aware negative selection. Stratified sampling guided by the positive set's taxonomic-group and IDP/non-IDP composition generated the final negative set (n = 14,813). Numbers indicate protein counts at each step.

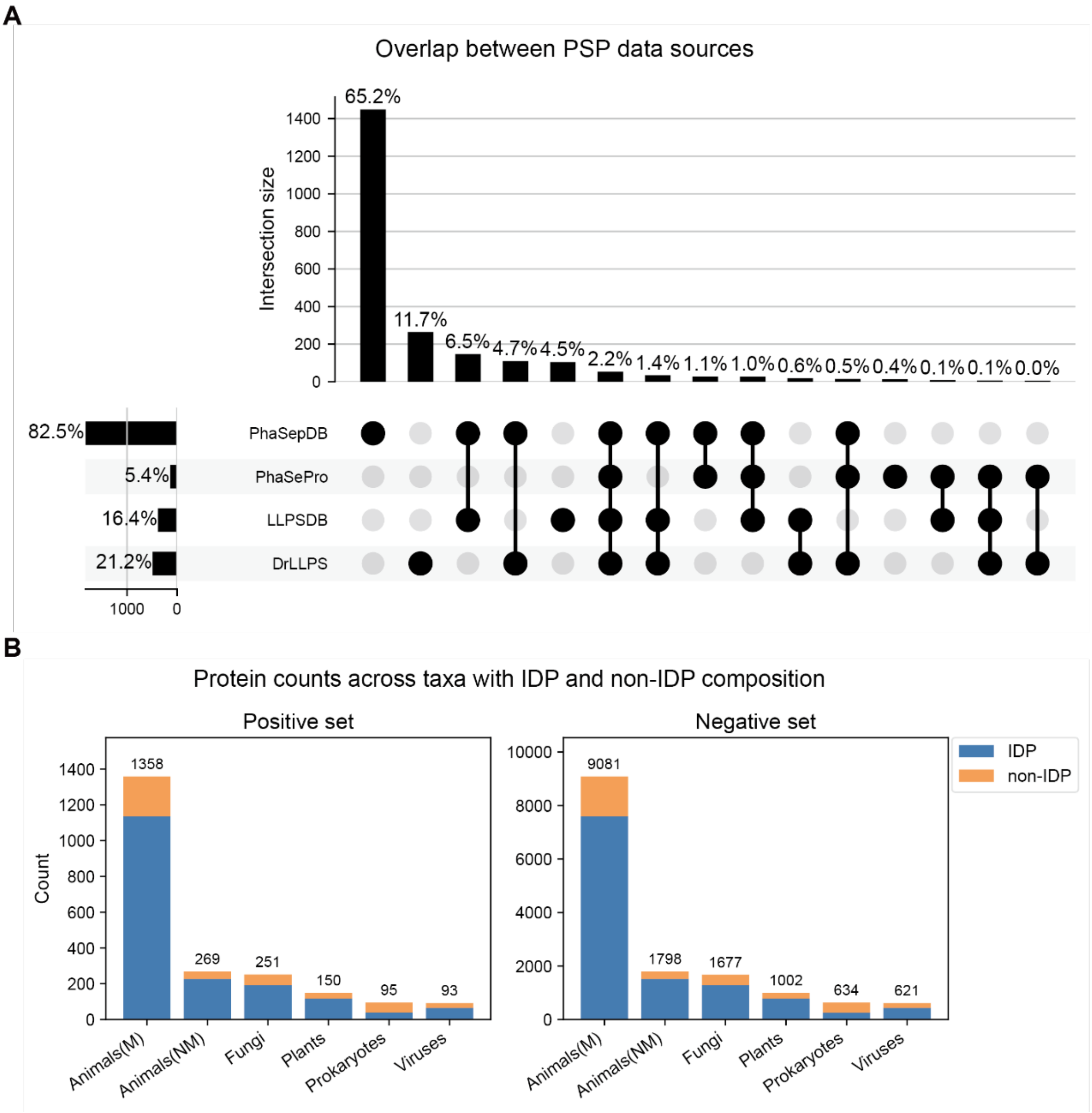


**Supplementary Figure 3**. Overview of taxonomy-aware benchmark dataset composition. **A** UpSet plot showing overlap among PSP records collected from four curated LLPS-focused resources (PhaSepDB, PhaSePro, LLPSDB and DrLLPS). Bars indicate the size of each intersection (top) and the total contribution of each resource (left). Filled dots denote the combination of resources defining each intersection; percentages report the fraction of the final positive set represented by each intersection. **B** Taxonomic-group composition of the benchmark positive and negative sets. Stacked bars show protein counts per taxonomic group (Animals(M), Animals(NM), Plants, Fungi, Prokaryotes and Viruses), partitioned into IDP and non-IDP subsets; numbers above bars indicate total counts per group.

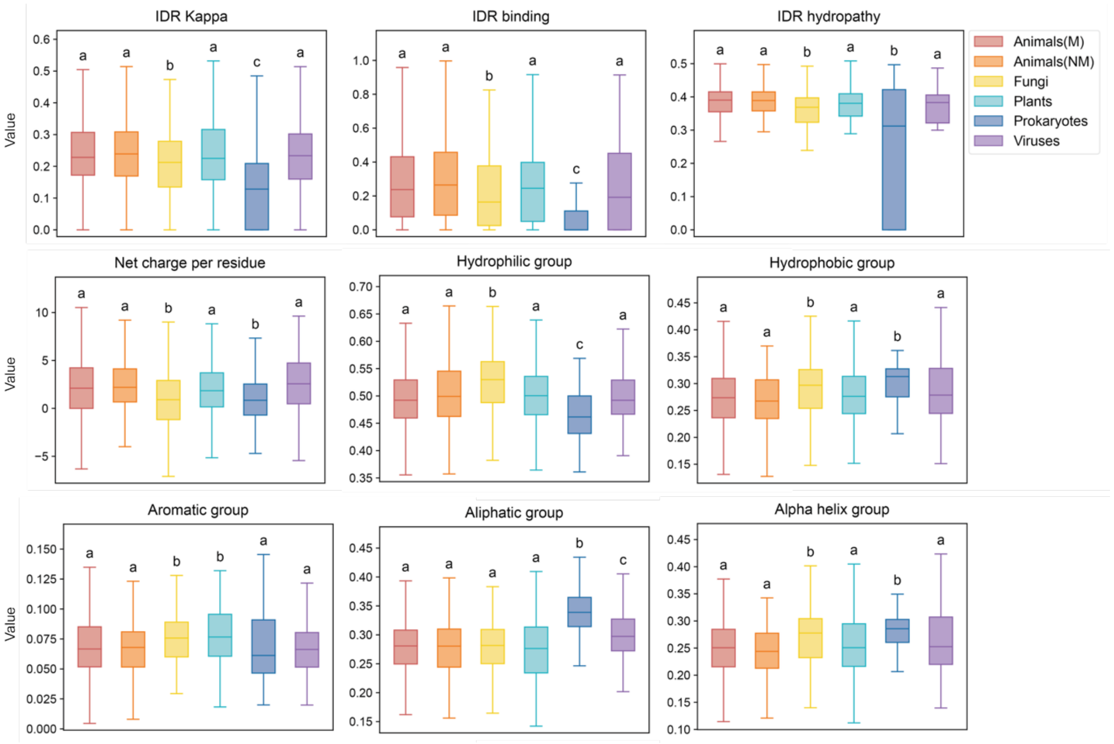


**Supplementary Figure 4**. Distributions of representative sequence and biophysical features for positive proteins across taxa, including IDR Kappa (κ; charge patterning), IDR binding score, IDR hydropathy, net charge per residue, and the fractions of residues belonging to hydrophilic (S, T, H, N, Q, E, D, K, R), hydrophobic (V, I, L, F, W, Y, M), aromatic (F, W, Y), aliphatic (A, V, L, I, M) and α-helix-promoting (V, I, Y, F, W, L) groups. Boxplots show the median (central line), interquartile range (IQR; 25th-75th percentiles), and whiskers extending to the most extreme values within 1.5 × IQR. Letters denote compact letter display (CLD) groupings derived from two-sided Mann-Whitney U tests with Benjamini-Hochberg false discovery rate (BH-FDR) correction; groups sharing at least one letter are not significantly different, whereas groups with no letters in common differ significantly (BH-FDR < 0.05).

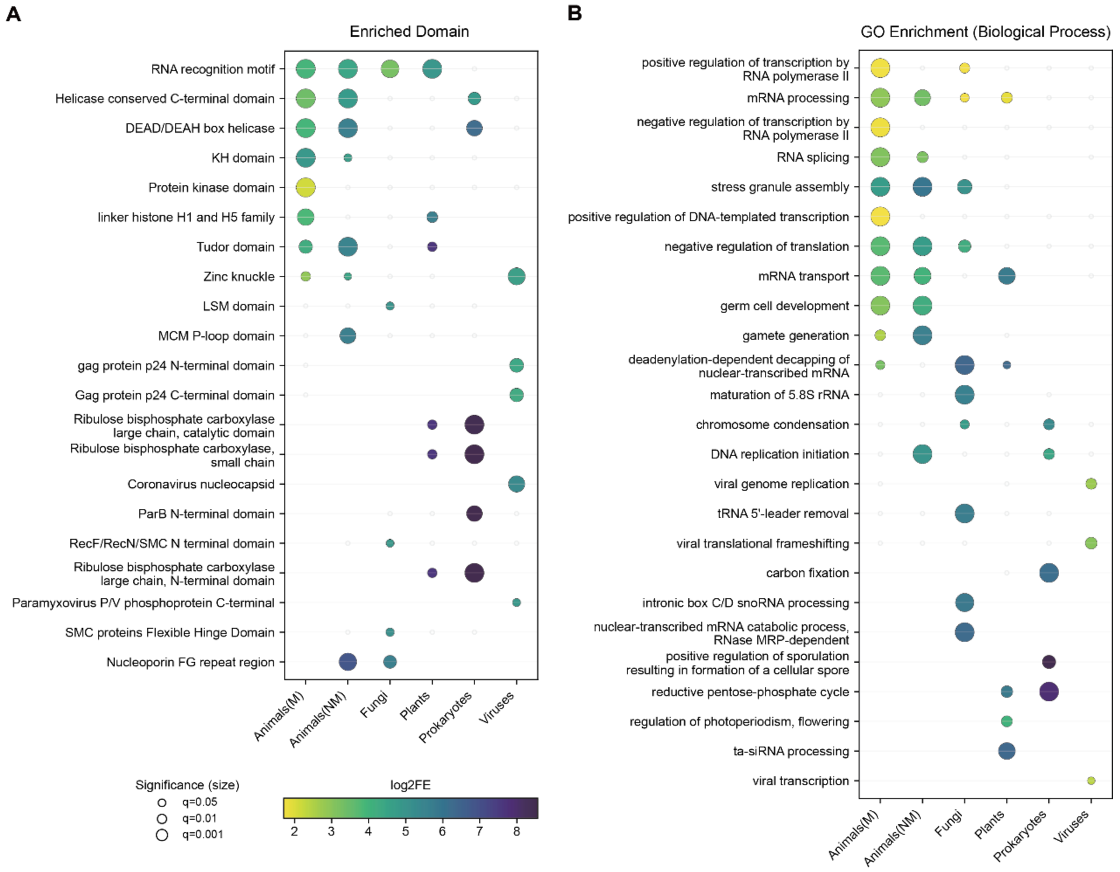


**Supplementary Figure 5**. Taxon-specific functional and domain enrichments of phase-separating proteins. Enrichment of Pfam domains (A) and Gene Ontology (GO) biological process terms (B) in positives across six taxonomic groups (Animals(M), Animals(NM), Plants, Fungi, Prokaryotes and Viruses). For each taxon, enrichment was calculated by comparing positives against the corresponding taxon-matched proteome background. Dot color encodes log2 fold enrichment (color bar), and dot size reflects enrichment significance (Benjamini-Hochberg FDR). Grey markers denote non-significant terms.

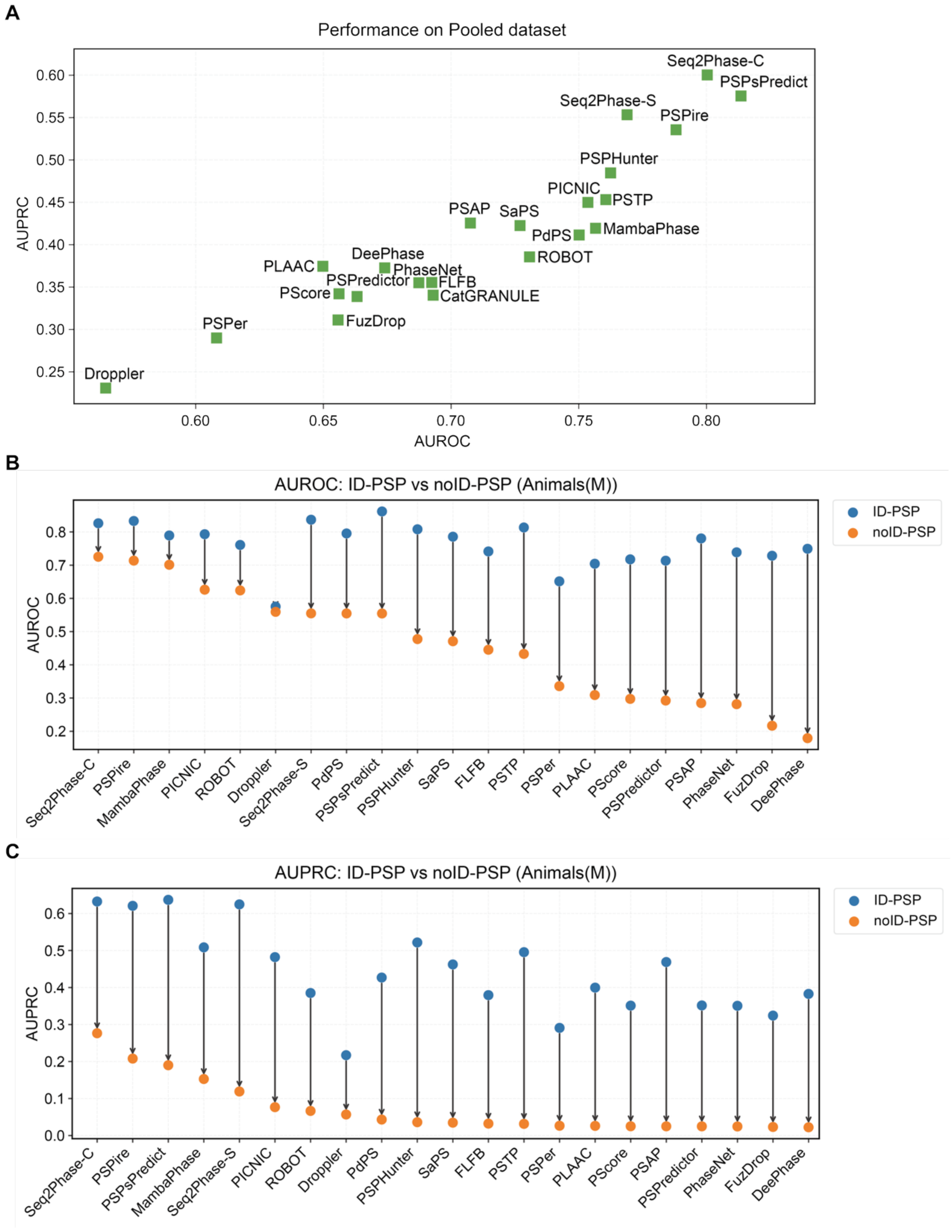


**Supplementary Figure 6.** Performance of predictors on the benchmark dataset. **A** Predictor performance on the pooled dataset combining Animals(M), Animals(NM), Plants and Fungi (green squares). Prokaryotes and Viruses were excluded because they are supported by only a subset of predictors, enabling a consistent comparison across all methods. **B and C** Predictor performance on phase-separating proteins (PSPs) with IDRs (ID-PSPs) and PSPs without IDRs (noID-PSPs) in the Animals(M) subset. Other subsets were not analyzed in this stratified manner because they contained too few noID-PSPs to support reliable evaluation. For each predictor, AUROC (B) and AUPRC (C) are reported separately for ID-PSPs (blue) and noID-PSPs (orange). Vertical connectors indicate the within-predictor performance gap between ID-PSP and noID-PSP subsets (arrow from ID-PSP to noID-PSPs).

#### SUPPLEMENTARY TABLES

Supplementary Table 1. List of proteins in benchmark positive and negative sets.
Supplementary Table 2. List of evaluated phase-separating protein predictors.
Supplementary Table 3. Predictor performance across six taxonomic groups in the benchmark dataset.
